# Copine-6 is a Ca^2+^ sensor for activity-induced AMPA receptor exocytosis

**DOI:** 10.1101/2023.05.10.540298

**Authors:** Jing Zhi Anson Tan, Se Eun Jang, Ana Batallas-Borja, Nishita Bhembre, Mintu Chandra, Lingrui Zhang, Huimin Guo, Mitchell T. Ringuet, Jocelyn Widagdo, Brett M. Collins, Victor Anggono

**Author notes:** These authors contributed equally to this work (Jing Zhi Anson Tan, Se Eun Jang). Present address: Department of Biological Sciences and Center for Structural Biology, Vanderbilt University, Nashville, TN 37232, USA.

## Abstract

The recruitment of synaptic AMPA (α-amino-3-hydroxy-5-methyl-4-isoxazole propionic acid) receptors underlies the strengthening of neuronal connectivity during learning and memory. This process is triggered by NMDA (*N*-methyl-*D*-aspartate) receptor-dependent postsynaptic Ca^2+^ influx. Synaptotagmin (Syt)-1 and −7 have been proposed as Ca^2+^-sensors for AMPA receptor exocytosis, but are functionally redundant. Here we identify a cytosolic C2 domain-containing Ca^2+^-binding protein Copine-6 that forms a complex with AMPA receptors. Loss of Copine-6 expression impairs activity-induced exocytosis of AMPA receptors in primary hippocampal neurons, which is rescued by wild-type Copine-6, but not Ca^2+^-binding mutants. In contrast, Copine-6 loss-of-function has no effects on steady-state expression or tetrodotoxin-induced synaptic upscaling of surface AMPA receptors. Loss of Syt-1/-7 significantly reduces Copine-6 protein expression. Interestingly, overexpression of wild-type Copine-6, but not the Ca^2+^-binding mutant, restores activity-dependent exocytosis of AMPA receptors in Syt-1/-7 double-knockdown neurons. We conclude that Copine-6 is a postsynaptic Ca^2+^-sensor that mediates AMPA receptor exocytosis during synaptic potentiation.

## INTRODUCTION

The ability of neurons to dynamically modulate their synaptic connection within neural networks provides a cellular basis for learning and memory (Kessels and Malinow, 2009; Takeuchi et al., 2014). *N*-methyl-*D*-aspartate (NMDA) receptor (NMDAR)-dependent long-term potentiation (LTP) is the most studied form of synaptic plasticity that plays an essential role in memory formation (Goto et al., 2021; Nabavi et al., 2014; Tsien et al., 1996). During LTP, activation of NMDARs mediates a large influx of calcium (Ca^2+^) into the postsynaptic compartment, thereby increasing the number of AMPA (α-amino-3-hydroxy-5-methyl-4-isoxazole propionic acid) receptors (AMPARs) at synapses and enhancing excitatory synaptic transmission (Anggono and Huganir, 2012; Hayashi, 2022; Huganir and Nicoll, 2013; Shi et al., 1999). The critical fusion of AMPAR-containing vesicles with the postsynaptic membrane is mediated by the soluble *N*-ethylmaleimide sensitive factor-attachment protein receptor (SNARE) complex (Jurado et al., 2013; Kennedy et al., 2010; Lledo et al., 1998). However, the mechanisms that underlie the Ca^2+^-dependent delivery of AMPARs to the plasma membrane remain unclear.

Two SNARE-associated proteins, Synaptotagmin-1 (Syt-1) and Synaptotagmin-7 (Syt-7), which bind Ca^2+^ with low and high affinity, respectively, have been proposed to act as postsynaptic Ca^2+^-sensors for AMPAR exocytosis during LTP (Wu et al., 2017). Given that LTP induction requires a brief increase of high postsynaptic Ca^2+^ concentration (Yang et al., 1999), selective activation of a low-affinity Ca^2+^-sensor, such as Syt-1, would in principle be ideal for mediating AMPAR exocytosis. However, the two Syts are functionally redundant as the re-expression of either Syt-1 or Syt-7 is sufficient to restore LTP-induced AMPAR exocytosis in Syt-1/-7 double-knockout neurons (Wu et al., 2017). Moreover, Syt-1 and Syt-7 are predominantly associated with synaptic vesicles and axonal plasma membranes, respectively (Sudhof, 2013; Vevea et al., 2021). The vastly different Ca^2+^-binding profiles of Syt-1 and Syt-7, as well as their expression and subcellular localization, suggest the possible existence of an alternative postsynaptic SNARE-associated Ca^2+^-binding protein that is essential for AMPAR exocytosis during synaptic potentiation.

## RESULTS

### Copine-6 binds Ca^2+^ via the C2B domain

In this study, we focused on a cytosolic Ca^2+^-and phospholipid-binding protein, Copine-6, which is highly enriched in the postsynaptic density (PSD) and is essential for hippocampal LTP, learning and memory (Burk et al., 2018; Reinhard et al., 2016). Copine-6 is highly expressed in excitatory neurons of the adult mouse hippocampus (Burk et al., 2018; Nakayama et al., 1999; Reinhard et al., 2016), and its level can be modulated by neuronal activity (Burk et al., 2018; Nakayama et al., 1998; Schiapparelli et al., 2022). The Copine family of proteins is characterized by two amino-terminal C2 domains and a carboxyl-terminal von Willebrand A (VWA) domain (Figure 1A). It is well-established that Copine-6 interacts with membranes and phospholipids in a Ca^2+^-dependent manner (Burk et al., 2018; Nakayama et al., 1999; Perestenko et al., 2010; Reinhard et al., 2016), but it is currently unknown if Copine-6 directly binds to Ca^2+^. To address this, we performed an isothermal titration calorimetry (ITC) assay which revealed that the full-length recombinant Copine-6 protein bound Ca^2+^ with a dissociation constant (K_d_) of 9.3 ± 1.8 μM (Figure 1B and Table S1). Mutations of acidic residues in the C2A domain (D93N/E95A, C2Amt) reduced Ca^2+^ binding by approximately 3-fold (K_d_ = 30.2 ± 3.1 μM, Figure 1C and Table S1), whereas those in the C2B domain (D229/231/237N, C2Bmt) reduced the affinity below detectable limits, suggesting that the C2B domain is the primary Ca^2+^ sensor (Figure 1D). We next determined the effects of these Copine-6 mutants on Ca^2+^-dependent membrane binding in cells. In HEK293T cells, the Ca^2+^ ionophore caused redistribution of HA-Copine-6 wild-type and C2Amt, but not C2Bmt, from cytosolic to the membrane fractions (Figure 1E). Similarly, Copine-6-GFP C2Bmt failed to translocate into the plasma membrane of rat primary hippocampal neurons following a brief stimulation with NMDA (Figure 1F and 1G). Taken together, these results suggest that Copine-6 directly binds to Ca^2+^ primarily through the C2B domain, which is essential for Copine-6 association with the plasma membrane in response to a rise in intracellular Ca^2+^ concentration.

**Figure 1.**
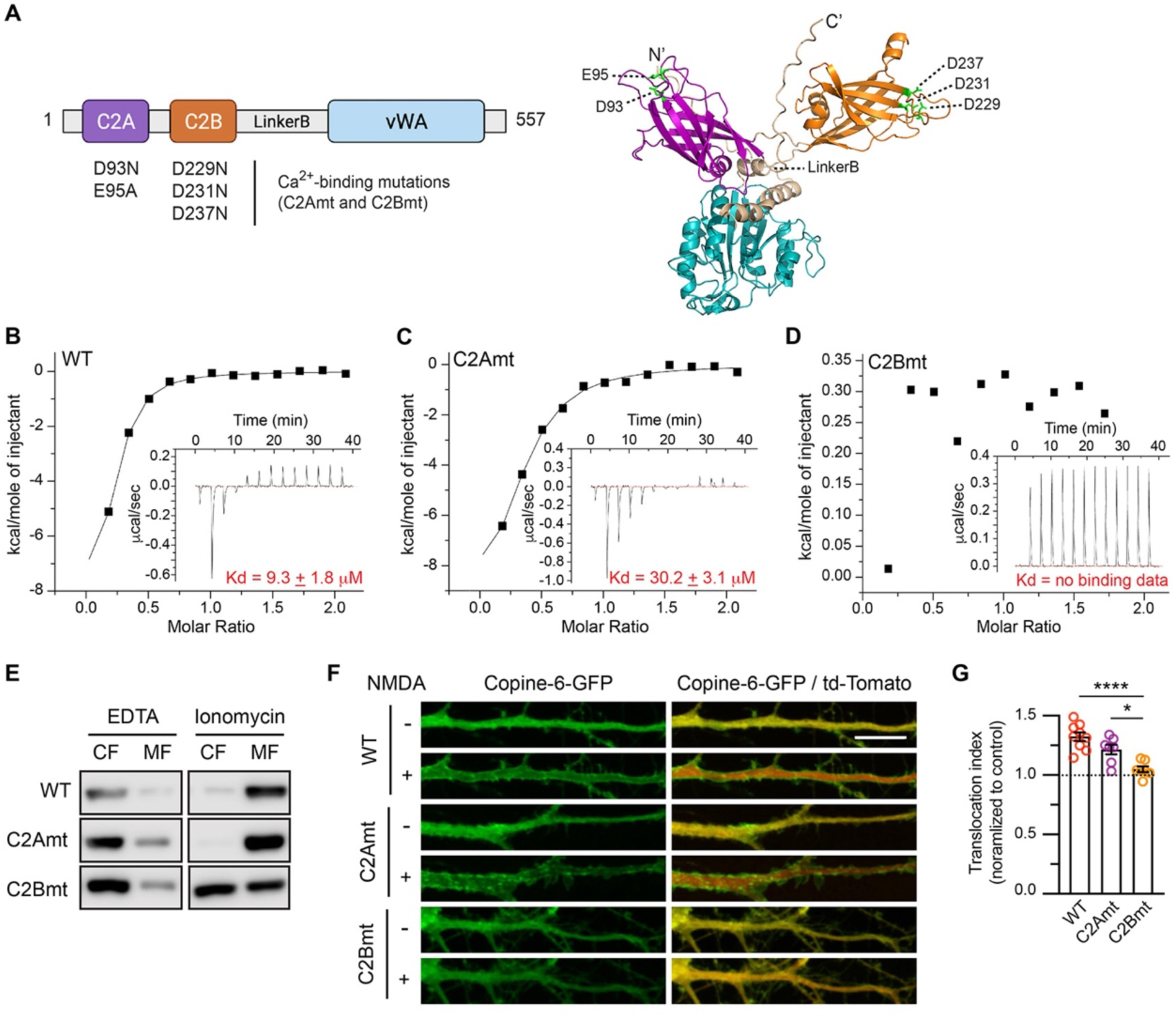
Direct binding of calcium to the C2B domain is essential for Copine-6 association with the plasma membrane. (A) Domain structure (left) and the AlphaFold predicted structure (right) of Copine-6, depicting the N-terminal C2A (magenta) and C2B (orange) Ca^2+^-binding domains, Linker-B region (beige) and vWA domain (cyan). The C2A (D93N and E95A) and C2B (D229N, D231N and D237N) Ca^2+^-binding mutants appear in green. (B–D) Isothermal titration calorimetry (ITC) experiments comparing the binding of 100 μM of full-length recombinant Copine-6 proteins, either (B) wild-type (WT), (C) D93N/E95Q (C2Amt) or (D) D229N/D231N/D237N (C2Bmt) mutants to 2 mM CaCl_2_ at 25°C. The thermodynamic parameters of Copine-6 binding to Ca^2+^ are shown in Table S1. (E) The translocation of Copine-6 to the membrane requires Ca^2+^-binding in the C2B domain. HEK293T cells were transfected with plasmids encoding HA-Copine-6 (WT, C2Amt or C2Bmt) for 48 h. Cells were treated with 2 mM EDTA or 2 μM ionomycin for 5 min, lyzed and subjected to ultracentrifugation. The presence of HA-Copine-6 in the cytosolic (CF) or membrane (MF) fractions was analyzed by immunoblotting with anti-HA antibodies. (F) Copine-6 C2Bmt fails to translocate to the neuronal membrane following NMDA stimulation. Hippocampal neurons were co-transfected with plasmids encoding td-Tomato (cytosolic marker) and Copine-6-GFP (WT, C2Amt, or C2Bmt) at DIV13. The translocation of Copine-6 from the cytosol into the plasma membrane was visualized by live-cell imaging in neurons treated with 10 μM NMDA and 100 μM glycine for 3 min in primary hippocampal neurons. Scale bar, 10 μm. (G) Quantification of the translocation index of Copine-6-GFP following NMDA treatment. Data represent mean ± SEM (*n* = 7-9 neurons per group from 3 independent experiments). \**P* < 0.05; \*\*\*\**P* < 0.0001 by using one-way ANOVA with Tukey’s multiple comparison test.

### Copine-6 interacts with the GluA1 subunit of AMPARs

To begin testing our hypothesis that Copine-6 is a postsynaptic Ca^2+^-sensor for activity-induced AMPAR exocytosis, we first determined whether Copine-6 interacts with AMPARs. Immunoprecipitation assays from hippocampal neuronal lysates revealed that the GluA1 and GluA2 subunits of AMPARs co-immunoprecipitate with anti-Copine-6 antibodies, but not by normal rabbit IgG (Figure 2A). Likewise, Copine-6 could be co-immunoprecipitated with anti-GluA1 antibodies (Figure 2A), confirming that it can form a complex with AMPARs in primary hippocampal neurons. We next determined whether Copine-6 can interact with the carboxyl-terminal (C-tail) of GluA1 or GluA2, which are crucial for regulating various aspects of AMPAR trafficking (Anggono and Huganir, 2012; Huganir and Nicoll, 2013). GST pull-down assays from HEK293T cell lysates showed that Myc-Copine-6 was pulled down specifically by GST-GluA1-C-tail but failed to interact with GST-GluA2-C-tail or free GST (Figure 2B). Pull-down assays with a series of GluA1 C-tail truncations (Figure 2C) mapped the binding region of Myc-Copine-6 to the membrane proximal region between amino acid 823 and 855 (Figure 2D and 2E). In addition, we revealed the Copine-6 C2 domains as the binding site for GluA1 C-tail (Figure 2F and 2G). Interestingly, deletion of the vWA domain, which is not required for GluA1 C-tail binding, markedly enhanced Copine-6 binding to GluA1 (Figure 2G), suggesting that the presence of this domain causes a steric hindrance and inhibits the binding of Copine-6 to its interacting partners.

**Figure 2.**
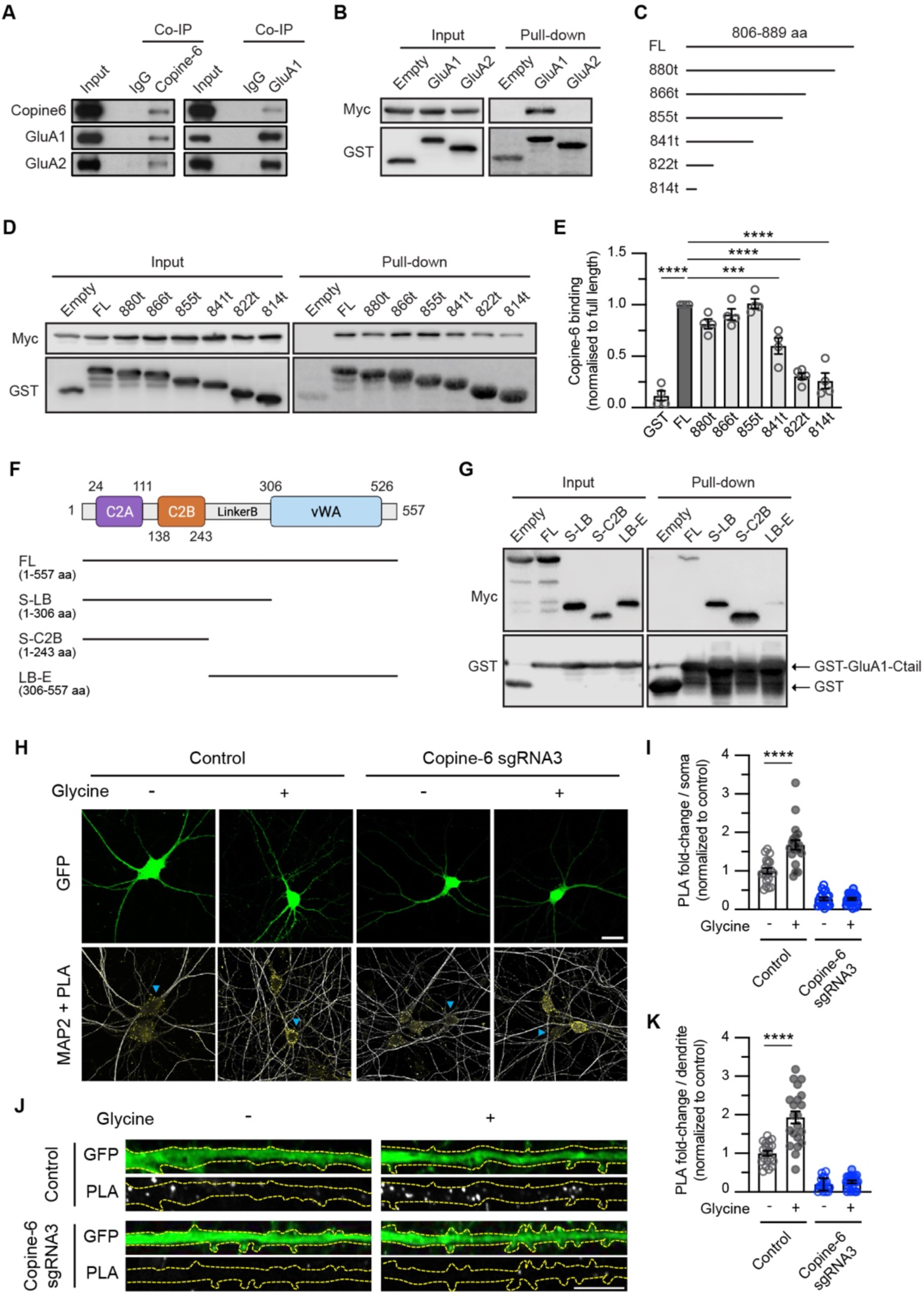
Glycine stimulation enhances the formation of the Copine-6 and AMPAR complex in hippocampal neurons. (A) Co-immunoprecipitation of Copine-6 and AMPARs. Lysates from primary neurons were immunoprecipitated (IP) with antibodies against Copine-6 and GluA1, or normal IgG (negative control). Total neuronal lysates (Input) and co-immunoprecipitated proteins were analyzed by immunoblotting with specific antibodies against Copine-6, GluA1 and GluA2. (B) HEK293T cells were co-transfected with plasmids encoding Myc-Copine-6 and either GST, GST-GluA1 C-tail or GST-GluA2 C-tail. Cells were lyzed and pulled down with GSH-Sepharose beads. Total lysates (Input) and bound proteins were resolved by SDS-PAGE and analyzed by western blotting with specific antibodies against Myc and GST. (C) Schematic diagram of GluA1 C-tail deletion constructs. (D) GST pull-down experiments from HEK293T cells co-expressing Myc-Copine-6 with GST alone or GST-GluA1 C-tails (full-length or truncated proteins). Total lysates and bound proteins were immunoblotted with anti-Myc and anti-GST antibodies. (E) Quantification of the levels of Copine-6 binding to GST-GluA1 full-length and truncated C-tails. Data represent mean ± SEM of band intensities normalized to GST-GluA1 C-tail full-length (*n* = 4). \*\*\**P* < 0.001; \*\*\*\**P* < 0.0001 using one-way ANOVA with Tukey’s multiple comparison test. (F) Schematic diagram of Myc-Copine-6 constructs used in the pull-down experiments. (G) HEK293T cell lysates co-expressing GST-GluA1 C-tail full-length and Myc-Copine-6 (either full-length or isolated domains). Total cell extracts and bound proteins were analyzed by immunoblotting with specific antibodies against Myc and GST. (H) Primary hippocampal neurons were transfected with plasmids encoding Cas9-GFP, either alone or with Copine-6 sgRNA3 at days in vitro (DIV) 12. At DIV 15, neurons were stimulated with glycine for 5 min and subjected to a proximity ligation assay (PLA) with GluA1 and Copine-6 antibodies. Representative images showing the PLA signals (yellow) and MAP2 staining (gray) in transfected hippocampal neurons (green, arrowheads). Scale bar, 20 μm. (I) Quantification of the number of PLA puncta in the soma. (J) Representative images showing the PLA puncta (gray) within 50 μm dendritic segments of transfected neurons (green). Scale bar, 10 μm. (K) Quantification of the number of PLA puncta in the dendrites. Data are normalized to the unstimulated control group and are represented as mean ± SEM (*n* = 19-21 neurons per group from 3 independent experiments). **** *P* < 0.0001 using one-way ANOVA with Tukey’s multiple comparison test.

To test the interaction between endogenous GluA1 and Copine-6, we performed proximity-based ligation assays (PLA) in primary hippocampal neurons under basal conditions and following glycine stimulation, a validated method of chemically inducing NMDAR-dependent LTP (chem-LTP) (Lu et al., 2001). GluA1-Copine-6 PLA puncta could be observed in both the soma and dendrites of neurons that expressed Cas9-GFP alone, which were significantly increased following glycine stimulation (Figure 2H–2K). In contrast, these PLA signals were significantly reduced in neurons expressing Cas9-GFP and a single-guide RNA (sgRNA) that specifically targets Copine-6 (Figure S1A and S1B) under both conditions, confirming the specificity of the PLA assay (Figure 2H–2K). Collectively, our data support the notion that the interaction between Copine-6 and GluA1 within the somatodendritic compartment is enhanced by LTP-inducing stimulation in neurons.

### Loss of Copine-6 impairs activity-induced AMPAR exocytosis

To examine the role of Copine-6 in AMPAR forward trafficking, we generated two independent sgRNAs, which effectively suppressed Copine-6 protein expression, but did not have any effect on the expression of Syt-1 and Syt-7 (Figure S1A and S1B). sgRNA-mediated knockdown of Copine-6 did not alter the steady-state expression of surface or total GluA1 measured with two complementary assays, namely immunostaining (Figure S1C and S1D) and surface biotinylation assays (Figure S1E–S1G). These data suggest that Copine-6 does not regulate the constitutive recycling of AMPARs under basal conditions. Our results also confirm that basal synaptic transmission is not altered in Copine-6-depleted neurons (Burk et al., 2018; Reinhard et al., 2016).

Using an antibody-feeding assay that binds to the extracellular domain of endogenous GluA1, we observed a significant increase in surface GluA1 expression in neurons expressing Cas9-GFP following glycine stimulation (Figure 3A and 3B). However, the chem-LTP-induced increase in surface GluA1 was impaired in neurons transfected with Cas9-GFP and Copine-6 sgRNA2 or sgRNA3 (Figure 3A and 3B). To corroborate our initial findings, we transduced primary cortical neurons with lentiviral particles expressing Cas9-GFP alone or Cas9-GFP with Copine-6 sgRNAs and performed surface biotinylation assays. As expected, chem-LTP stimulation significantly enhanced GluA1 expression on the plasma membrane of neurons expressing Cas9-GFP alone (Figure 3C and 3D). By contrast, neurons expressing Cas9-GFP and Copine-6 sgRNA2 or sgRNA3 failed to exhibit upregulation of surface GluA1 following glycine treatment (Figure 3C and 3D). Finally, to directly determine the role of Copine-6 in AMPAR exocytosis, we visualized the insertion of AMPARs by imaging super-ecliptic pH-sensitive-GFP (SEP)-tagged GluA1 with total internal reflection fluorescence (TIRF) microscopy in live neurons (Lin et al., 2009). In control neurons expressing Cas9-mCherry alone, there was a significant increase in the number of SEP-GluA1 insertions on the dendritic plasma membrane following chem-LTP (Figure 3E and 3F). Glycine-induced enhancement of GluA1 insertion events was abolished in neurons expressing Cas9-mCherry with Copine-6 sgRNA3 (Figure 3E and 3F). Together, our results demonstrate that postsynaptic Copine-6 deficiency impairs AMPAR exocytosis during synaptic potentiation in neurons, despite normal expression of both Syt-1 and Syt-7.

**Figure. 3.**
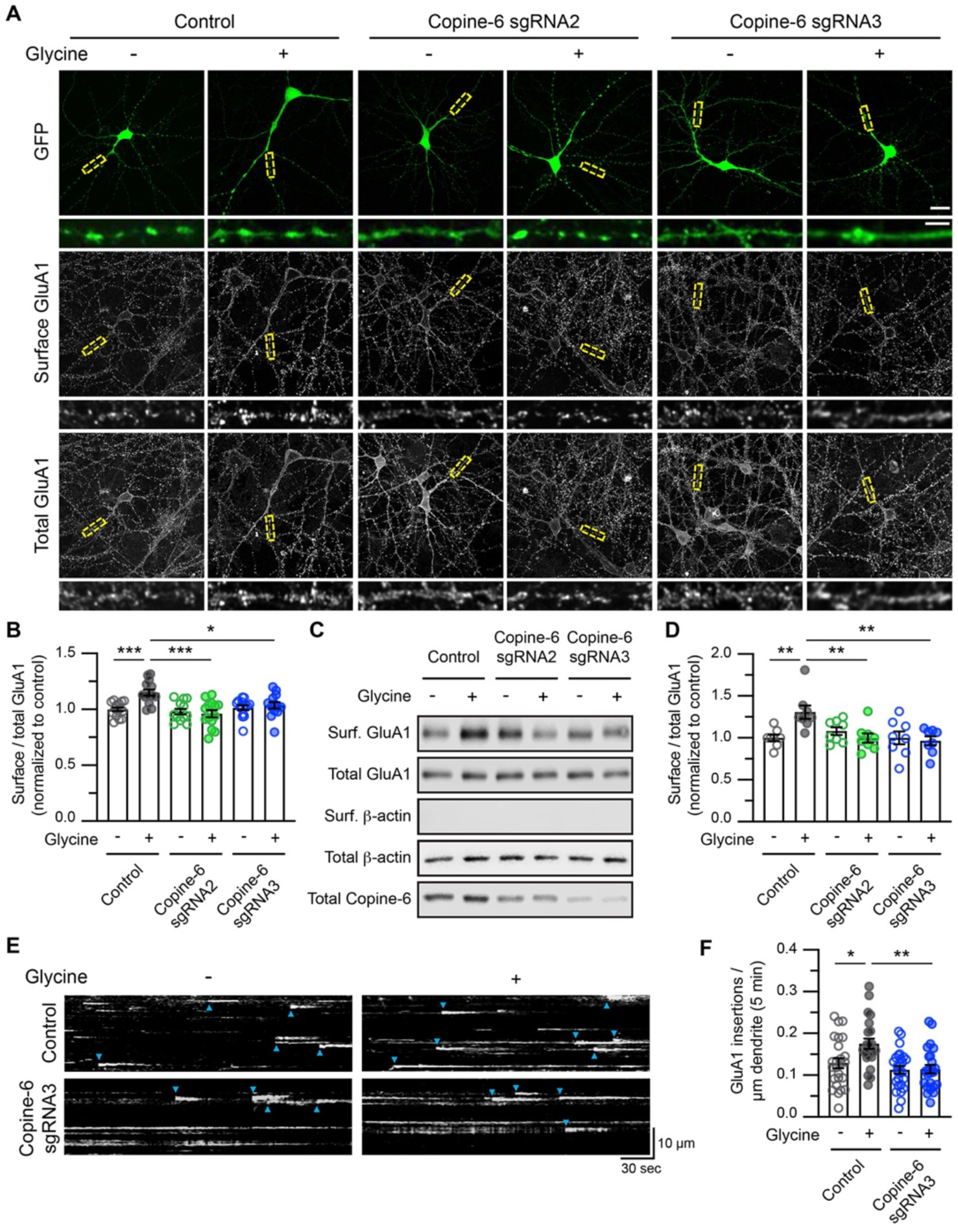
Loss of Copine-6 expression impairs the glycine-induced exocytosis of AMPARs. (A) Hippocampal neurons expressing Cas9-GFP, either alone or with Copine-6 sgRNA2 or sgRNA3 were stimulated with glycine for 5 min. Live neurons were incubated with a specific antibody against the N-terminus of GluA1 at 4°C (surface GluA1), followed by fixation, permeabilization and staining with a specific antibody against the C-terminus of GluA1 (total GluA1). Representative images of the surface and total GluA1 staining in neurons from each group are shown (top panels). Scale bar, 20 μm. The bottom panels show enlarged images of dendritic segments within the boxed regions. Scale bar, 5 μm. (B) Quantification of the surface/total GluA1 ratio normalized to non-stimulated control neurons (*n* = 13-15 neurons per group from 3 independent experiments). (C) Primary cortical neurons were transduced with lentiviral particles expressing Cas9-GFP, either alone or with Copine-6 sgRNA2 or sgRNA3 at DIV 9. Transduced neurons were stimulated with glycine for 5 min and subjected to a surface biotinylation assay at DIV 15. The relative amounts of surface and total proteins were assessed by western blotting using specific antibodies against GluA1, Copine-6 and β-actin. (D) Quantification of the surface/total GluA1 ratio in Copine-6-depleted neurons following glycine stimulation, normalized to non-stimulated control neurons (*n* = 8 cultures per group from 3 independent experiments). (E) Hippocampal neurons were co-transfected with plasmids encoding SEP-GluA1 and Cas9-mCherry control or Cas9-mCherry and Copine-6 sgRNA3. SEP-GluA1 insertion events over a 5 min period in non-stimulated and glycine-stimulated neurons were visualized using a TIRF microscope. Representative *y*-axis by time (*y*-t) maximum intensity projections of GluA1 insertion events in a 30 μm segment of a secondary dendrite from a neuron in each group. Each GluA1 insertion event is marked by a blue arrowhead. (F) Quantification of the GluA1 insertion events per μm of dendrite over 5 min following glycine stimulation (*n* = 23-26 dendritic segments from 4 independent experiments). All data are represented as mean ± SEM. \**P* < 0.05; \*\**P* < 0.01; \*\*\**P* < 0.001 using one-way ANOVA with a Tukey’s multiple comparison test.

To ascertain that Copine-6 plays a specific role in chem-LTP-induced AMPAR exocytosis, we next assessed the effect of Copine-6 knockdown using a homeostatic synaptic scaling paradigm induced by chronic synaptic silencing (Anggono et al., 2011). Interestingly, a 48-h treatment with tetrodotoxin (TTX) significantly reduced Copine-6 expression by ∼40% (Figure S2A and S2B), consistent with previous findings that an increase in neuronal activity can upregulate Copine-6 expression (Burk et al., 2018; Nakayama et al., 1998; Schiapparelli et al., 2022). Despite the reduction in Copine-6 expression, the level of surface GluA1 expression was robustly upregulated in control neurons following chronic neuronal inactivity (Figure S2A and S2C). It is therefore not surprising that Copine-6 knockdown had no effect on TTX-induced AMPAR recruitment onto the plasma membrane (Figure S2A and S2C). Two studies have also shown that Copine-6 is not required for NMDAR-dependent long-term depression (LTD) (Burk et al., 2018; Reinhard et al., 2016). Thus, Copine-6 plays a very specific role in mediating the exocytosis of AMPARs during NMDAR-dependent LTP, and is not involved in homeostatic synaptic up-scaling or LTD.

### Copine-6 binding to Ca^2+^ is essential for activity-induced AMPAR exocytosis

To interrogate the functional significance of Ca^2+^-binding on Copine-6 in regulating the trafficking of AMPARs, we performed rescue experiments in Copine-6-depleted neurons by re-expressing sgRNA-resistant HA-Copine-6, either wild-type, C2Amt or C2Bmt. Expression of Copine-6 wild-type fully restored the chem-LTP deficit in neurons expressing Cas9-GFP and Copine-6-sgRNA3 using three complementary assays, namely the antibody-feeding assay (Figure 4A and 4B), surface biotinylation assay (Figure 4C and 4D) and TIRF microscopy imaging of SEP-GluA1 insertions (Figure 4E and 4F). By contrast, both Copine-6 C2Amt and C2Bmt failed to support the glycine-induced insertion of GluA1 to the neuronal plasma membrane (Figure 4). Importantly, overexpression of Copine-6, either wild-type or C2 mutants, did not affect the steady-state expression of surface and total GluA1 in neurons, suggesting that Copine-6 mutants do not exert dominant-negative effects under basal conditions (Figure S3).

**Figure 4.**
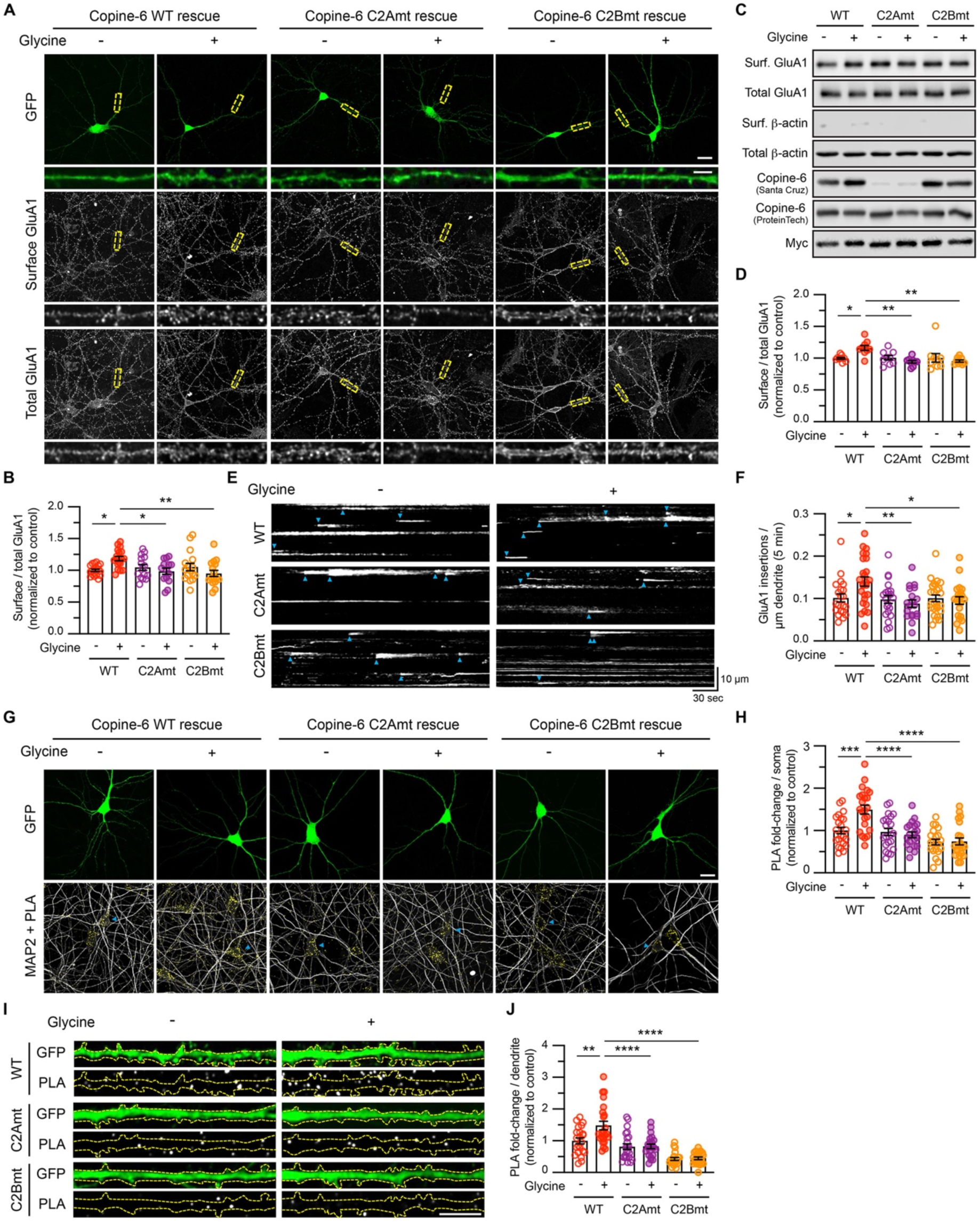
Copine-6 binding to Ca^2+^ is required for glycine-induced exocytosis of AMPARs. (A) Copine-6-depleted hippocampal neurons (expressing Cas9-GFP and Copine-6 sgRNA3) were transfected with plasmids encoding sgRNA3-resistant HA-Copine-6 cDNAs, either wild-type (WT), C2Amt (D93N/E95A) or C2Bmt (D229/231/237N) at DIV 12. At DIV 15, live neurons were stimulated with glycine for 5 min, and incubated with a specific antibody against the N-terminus of GluA1 at 4°C (surface GluA1). They were then fixed, permeabilized and stained with a specific antibody against the C-terminus of GluA1 (total GluA1). Representative images of the surface and total GluA1 staining in neurons from each group are shown (top panels). Scale bar, 20 μm. The bottom panels show enlarged images of dendritic segments within the boxed regions. Scale bar, 5 μm. (B) Quantification of the surface/total GluA1 ratio normalized to non-stimulated HA-Copine-6 WT neurons (*n* = 14-18 neurons per group from 3 independent experiments). (C) Primary cortical neurons were transduced with lentiviral particles expressing Cas9-GFP and Copine-6 sgRNA3, and sgRNA3-resistant Myc-Copine-6, either WT, C2Amt or C2Bmt at DIV9. Transduced neurons were stimulated with glycine for 5 min and subjected to a surface biotinylation assay at DIV15. The relative amounts of surface and total proteins were assessed by western blotting using specific antibodies against GluA1, Copine-6, Myc and β-actin. The Santa Cruz antibody failed to recognize Copine-6 C2Amt, revealing the extent of endogenous Copine-6 depletion in these neurons. (D) Quantification of the surface/total GluA1 ratio in Copine-6 rescued neurons following glycine stimulation, normalized to non-stimulated Myc-Copine-6 WT neurons (*n* = 9 cultures per group from 5 independent experiments). (E) Hippocampal neurons were co-transfected with plasmids encoding SEP-GluA1, Cas9-mCherry and Copine-6 sgRNA3, together with HA-Copine-6 sgRNA3-resistant constructs, either WT, C2Amt or C2Bmt. SEP-GluA1 insertion events over a 5 min period in non-stimulated and glycine-stimulated neurons were visualized using a TIRF microscope. Representative *y*-axis by time (*y*-t) maximum intensity projections of GluA1 insertion events in a 30 μm segment of a secondary dendrite from a neuron in each group. Each GluA1 insertion event is marked by a blue arrowhead. (F) Quantification of the GluA1 insertion events per μm of dendrite over 5 min following glycine stimulation (*n* = 19-26 dendritic segments from 4 independent experiments). (G) Primary hippocampal neurons were co-transfected with plasmids encoding Cas9-GFP and Copine-6 sgRNA3, together with HA-Copine-6 sgRNA3-resistant constructs (WT, C2Amt or C2Bmt) at DIV 12. At DIV 15, neurons were stimulated with glycine for 5 min and subjected to a proximity ligation assay (PLA) with GluA1 and Copine-6 antibodies. Representative images showing the PLA signals (yellow) and MAP2 staining (gray) in transfected hippocampal neurons (green, arrowheads). Scale bar, 20 μm. (H) Quantification of the number of PLA puncta in the soma. (I) Representative images showing the PLA puncta (gray) within 50 μm dendritic segments of transfected neurons (green). Scale bar, 10 μm. (J) Quantification of the number of PLA puncta in the dendrites. Data are normalized to the unstimulated HA-Copine-6 WT group (*n* = 21-23 neurons per group from 4 independent experiments). All data are represented as mean ± SEM. \**P* < 0.05; \*\**P* < 0.01; \*\*\**P* < 0.001; **** *P* < 0.0001 using one-way ANOVA with a Tukey’s multiple comparison test.

Although our C2B mutant data demonstrate that Copine-6 binding to Ca^2+^ is critical for chem-LTP-induced AMPAR exocytosis, we were intrigued by the effects produced by C2Amt, which can still bind to Ca^2+^ (albeit with a 3-fold reduction in Ca^2+^-binding affinity) and translocate to the plasma membrane in response to Ca^2+^ influx. Because GluA1 C-tail interacts with Copine-6 through its C2 domains, we asked whether the association of Copine-6 with GluA1 in response to glycine stimulation was affected by mutations in the C2 domains. In Copine-6-depleted neurons, overexpression of HA-Copine-6 wild-type restored the activity-dependent augmentation in the interaction with endogenous GluA1 as shown by an increase in the number of PLA puncta in the soma (Figure 4G and 4H) and dendrites (Figure 4I and 4J) of primary hippocampal neurons. Both C2Amt and C2Bmt robustly inhibited the glycine-induced increase in the interaction between Copine-6 and GluA1 (Figure 4G–4J). Furthermore, C2Bmt exhibited a significantly lower number of PLA puncta under basal conditions, demonstrating that the binding of Copine-6 to Ca^2+^ is critical for its interaction with GluA1, a property that can be modulated by chem-LTP stimulation. These results also suggest that the association with GluA1 is sensitive to alterations in Copine-6 binding affinity to Ca^2+^, and thus its conformational change, or that Asp-93 and/or Glu-95 may be directly involved in GluA1 binding, providing an explanation for the apparent effect of C2Amt on glycine-induced AMPAR exocytosis.

As the major Ca^2+^-sensors of the synchronous and asynchronous release of neurotransmitters, the loss of Syt-1 and Syt-7 severely impairs the synaptic vesicle exocytosis evoked by action potential firing or trains of high-frequency stimuli in mammalian central neurons (Bacaj et al., 2013; Wu et al., 2017). The fact that Copine-6 expression is sensitive to action potential blockade (Figure S2) prompted us to investigate the level of Copine-6 protein in Syt-1/Syt-7 double knockdown neurons. Consistent with previous findings, simultaneous depletion of Syt-1 and Syt-7 blocked glycine-induced insertion of GluA1 to the plasma membrane, and as expected, it also caused a significant downregulation in the level of the Copine-6 protein expression (Figure S4A and 4B). Importantly, overexpression of HA-Copine-6 wild-type, but not C2Bmt, completely restored the chem-LTP-induced increase in the level of surface GluA1 expression in Syt-1/Syt-7 double knockdown neurons (Figure S4A and S4C). Thus, the reduction of Copine-6 expression may account for the apparent impairment of activity-dependent exocytosis of AMPARs in the absence of Syt-1 and Syt-7. Unlike the proximal interaction between GluA1 and Copine-6, we barely detect any PLA signals using antibodies against Syt-1 and GluA1 (Figure S4D and S4E). These results further indicate that Syt-1 and GluA1 may not exist within the same subcellular compartment under basal conditions or following glycine stimulation.

## DISCUSSION

The recruitment of AMPARs at synapses generally involves three distinct steps: exocytosis at extra- or peri-synaptic sites, lateral diffusion to synapses, and trapping by scaffolding molecules at the PSD (Makino and Malinow, 2009; Opazo and Choquet, 2011; Penn et al., 2017). The activity-dependent insertion of AMPARs is essential for maintaining LTP as it provides a continuous supply of and replenishes synaptic receptors. Here we provide several lines of evidence that directly support a role for Copine-6 as the postsynaptic Ca^2+^-sensor that mediates AMPAR exocytosis during synaptic potentiation. Firstly, Copine-6 interacts with the C-tail of the GluA1 subunit of AMPARs. The association between Copine-6 and GluA1 is enhanced by chem-LTP stimulation, which requires direct Ca^2+^ binding to the Copine-6 C2B domain. Secondly, loss of Copine-6 expression selectively impairs the insertion of AMPARs into the plasma membrane during NMDAR-dependent chem-LTP but is not required for TTX-induced homeostatic synaptic upscaling of AMPARs. In addition, modulation of Copine-6 expression has no effect on the steady-state levels of AMPARs under basal conditions. Thirdly, activity-induced exocytosis of AMPARs requires the direct binding of Ca^2+^ to the C2B domain of Copine-6 and activity-dependent association with AMPARs. Finally, overexpression of wild-type Copine-6, but not the Ca^2+^-binding defective C2B mutant, restores the activity-induced insertion of AMPARs in Syt-1/Syt-7 double knockdown neurons. However, the possibility that Copine-6 and Syt-1/-7 have sequential roles in mediating AMPAR exocytosis cannot be ruled out at present.

Our conclusion is also supported by several molecular features of Copine-6 that make it a suitable postsynaptic Ca^2+^-sensor that mediates AMPAR exocytosis during LTP. Copine-6 is highly enriched in the somatodendritic compartment and PSD (Burk et al., 2018; Nakayama et al., 1999; Reinhard et al., 2016), where AMPARs are localized. It binds to Ca^2+^ with a moderate affinity, which subsequently triggers its interaction with phospholipids and translocation to vesicles or endosomes, the plasma membrane and dendritic spines (Burk et al., 2018; Nakayama et al., 1999; Perestenko et al., 2010; Reinhard et al., 2016). Importantly, Copine-6 also binds to components of the SNARE complex in a Ca^2+^-dependent manner (Liu et al., 2018), including synaptobrevin-2 and syntaxin-4, which are required for activity-dependent exocytosis of AMPARs during LTP (Jurado et al., 2013; Kennedy et al., 2010). Moreover, Copine-6 expression itself can be upregulated in response to heightened neuronal activity to support the increased demand for AMPAR exocytosis during LTP (Burk et al., 2018; Nakayama et al., 1998; Reinhard et al., 2016).

In addition to its role in mediating AMPAR exocytosis during synaptic potentiation, Copine-6 also plays an important role in structural plasticity. It interacts with the small GTPase Rac1 and regulates the dynamics of the actin cytoskeleton, a process that is essential for activity-dependent spine enlargement during LTP (Reinhard et al., 2016). It also mediates brain-derived neurotrophic factor (BDNF)-induced formation of mushroom spines by controlling the recycling of BDNF receptors (TrkB) in hippocampal neurons (Burk et al., 2018). Our discovery of the essential role of postsynaptic Copine-6 as a Ca^2+^ sensor for AMPAR exocytosis during synaptic potentiation not only provides new insight into the mechanism of synaptic plasticity but also positions Copine-6 as a versatile Ca^2+^ sensor that integrates postsynaptic NMDAR-dependent Ca^2+^ signaling that supports both functional and structural plasticity during LTP.

## ACKNOWLEDGEMENTS

We thank R. Amor for support with microscopy; R. Tweedale for comments on the manuscript; and P. Sah and J. Götz for insightful discussions. This work was supported by Australian National Health and Medical Research Council (NHMRC) Project Grant GNT1138452 (to V.A. and B.M.C.), Australian Research Council (ARC) Discovery Project Grant DP220101645 (to V.A.) and Australian Medical Research Future Fund (Clem Jones Centre for Ageing Dementia Research Flagship Project Grant to V.A.). V.A. holds an ARC Future Fellowship (FT220100485). J.Z.A.T. was supported by a University of Queensland (UQ) Researcher Retention (RSA2) Fellowship. J.W. was supported by a UQ Amplify award. S.E.J., A.B.B. and L.Z. were supported by UQ Research Training Scholarships. Imaging was performed at the Queensland Brain Institute’s Advanced Microscopy Facility, supported by the Australian Government through ARC LIEF grant LE130100078.

## AUTHOR CONTRIBUTIONS

Conceptualization: J.Z.A.T., S.E.J., B.M.C. and V.A. Methodology: J.Z.A.T, S.E.J., N.B., M.C., A.B.B., L.Z., H.G., M.T.R., J.W., B.M.C. and V.A. Investigation: J.Z.A.T, S.E.J., N.B., M.C., A.B.B., L.Z., H.G., M.T.R., J.W., B.M.C. and V.A. Visualization: J.Z.A.T. and V.A. Funding acquisition: B.M.C. and V.A. Project administration: V.A. Supervision: B.M.C. and V.A. Writing – original draft: J.Z.A.T., S.E.J., and V.A. Writing – review & editing: J.Z.A.T., M.C., J.W., B.M.C. and V.A.

## STAR METHODS

### KEY RESOURCES TABLES

| REAGENT or RESOURCE | SOURCE | IDENTIFIER |
| --- | --- | --- |
| **Antibodies** |  |  |
| Alexa 568-conjugated goat anti-rabbit IgG antibody | Thermo Scientific | Cat# A-11036<br>RRID: AB_10563566 |
| Alexa 647-conjugated goat anti-chicken IgY antibody | Abcam | Cat# ab150175;<br>RRID: AB_2732800 |
| Alexa 647-conjugated goat anti-rabbit IgG antibody | Thermo Scientific | Cat# A-21245<br>RRID: AB_2535813 |
| β-actin (mouse monoclonal, clone C4, used 1:5,000 in WB) | Santa Cruz Biotechnology | Cat# sc-47778,<br>RRID: AB_2714189 |
| Copine-6 (mouse monoclonal, clone 42, used 1:500 in WB) | Santa Cruz Biotechnology | Cat# sc-136357;<br>RRID: AB_10610096 |
| Copine-6 (rabbit polyclonal, used 1:200 in ICC and PLA, 1:1,000 in WB and in IP) | ProteinTech | Cat# 13782-1-AP;<br>RRID: AB_2292172 |
| GluA1 C-terminal (mouse monoclonal, clone 160E5, used 1:200 in PLA) | Synaptic Systems | Cat# 182011; RRID: AB_2113443 |
| GluA1 C-terminal (rabbit polyclonal, used 1:100 in ICC and 1:1000 in WB) | Merck Millipore | Cat# ab1504; RRID: AB_2113602 |
| GluA1 N-terminal (rabbit polyclonal, used 1:500 in ICC) | Merck Millipore | Cat# abn241; RRID: AB_2721164 |
| GluA1 N-terminal (mouse monoclonal, clone RH95, used in IP) | Merck Millipore | Cat# MAB2263;<br>RRID: AB_11212678 |
| GluA2 (rabbit polyclonal, used 1:2,000 in WB) | Merck Millipore | Cat# 07-598; RRID: AB_310741 |
| GST (mouse monoclonal, clone 3G12B10, used 1:5,000 in WB) | Proteintech | Cat# 66001-2-Ig |
| MAP2 (chicken polyclonal, used 1:2,000 in ICC) | Abcam | Cat# ab92434;<br>RRID: AB_2138147 |
| Myc (mouse monoclonal, clone 9E10, used 1:5,000 in WB) | Santa Cruz Biotechnology | Cat# sc-40,<br>RRID: AB_627268 |
| Synaptotagmin-1 (mouse monoclonal, clone 41.1, used 1:1,000 in WB and 1:200 in PLA) | Synaptic Systems | Cat# 105011, RRID: AB_887832 |
| Synaptotagmin-7 (rabbit polyclonal, used 1:1,000 in WB) | Synaptic Systems | Cat# 105173, RRID: AB_887838 |
| Peroxidase-conjugated donkey anti-rabbit IgG antibody | GE Healthcare | Cat# NA934,<br>RRID: AB_772206 |
| Peroxidase-conjugated sheep anti-mouse IgG antibody | GE Healthcare | Cat# NA931,<br>RRID: AB_772210 |
| **Chemicals, Peptides, and Recombinant Proteins** |  |  |
| Bicuculline methiodide | Abcam | Cat# ab120108 |
| Can Get Signal Immunoreaction Enhancer Solution | Toyobo Life Science | Cat# NKB-601 |
| cOmplete, EDTA-free protease inhibitor cocktail | Sigma-Aldrich | Cat# 11836170001 |
| Duolink in situ PLA probe anti-mouse PLUS | Sigma-Aldrich | Cat# DUO92001 |
| Duolink in situ detection reagents red | Sigma-Aldrich | Cat# DUO92008 |
| Glutathione agarose | Thermo Scientific | Cat# 16100 |
| Lipofectamine 2000 | Thermo Scientific | Cat# 11668019 |
| Neutravidin agarose | Thermo Scientific | Cat# 29204 |
| Papain suspension | Worthington | Cat# LS003126 |
| Strychnine hydrochloride | Sigma | Cat# S8753 |
| Sulfo-NHS-SS-biotin | CovaChem | Cat# 14207-100 |
| Tetrodotoxin citrate | Abcam | Cat# ab120055 |
| **Experimental Models: Cell Lines** |  |  |
| Human: HEK293T | ATCC | Cat# CRL-3216,<br>RRID:CVCL_0063 |
| **Experimental Models: Organisms/Strains** |  |  |
| Rat: Sprague-Dawley | University of<br>Queensland Biological<br>Resources | RRID:RGD_70508 |
| **Recombinant DNA** |  |  |
| FUW-Copine-6-sgRNA_resistant<br>(wild-type, C2Amt and C2Bmt) | This paper | N/A |
| FG12-Syt-1/Syt-7-shRNA | This paper | N/A |
| pCAG-SEP-GluA1 | R. Huganir (Johns<br>Hopkins University) | N/A |
| pCIS | R. Huganir (Johns<br>Hopkins University) | N/A |
| pCIS-GluA1-C-tail full-length (aa 806-889) | This paper | N/A |
| pCIS-GluA1-C-tail 880t (aa 806-880) | This paper | N/A |
| pCIS-GluA1-C-tail 866t (aa 806-866) | This paper | N/A |
| pCIS-GluA1-C-tail 855t (aa 806-855) | This paper | N/A |
| pCIS-GluA1-C-tail 841t (aa 806-841) | This paper | N/A |
| pCIS-GluA1-C-tail 822t (aa 806-822) | This paper | N/A |
| pCIS-GluA1-C-tail 814t (aa 806-814) | This paper | N/A |
| pCIS-GluA2-C-tail full-length (aa 833-882) | This paper | N/A |
| pEGFP-N1 | Clontech | N/A |
| pEGFP-N1-Copine-6-sgRNA_resistant<br>(wild-type, C2Amt and C2Bmt) | This paper | N/A |
| pHA-N1 | This paper | N/A |
| pHA-N1-Copine-6-sgRNA_resistant<br>(wild-type, C2Amt and C2Bmt) | This paper | N/A |
| pLentiCRISPR-GFP | S. Incontro<br>(Incontro et al., 2018) | N/A |
| pLentiCRISPR-GFP-Copine-6-sgRNA2 | This paper | N/A |
| pLentiCRISPR-GFP-Copine-6-sgRNA3 | This paper | N/A |
| pMD2.G | Addgene (D. Trono) | Plasmid #12259 |
| pMDLg/pRRE | Addgene (D. Trono) | Plasmid #12251 |
| pRK5-myc-Copine-6-Full-length (1-557aa) | This paper | N/A |
| pRK5-myc-Copine-6-Start-LinkerB (1-306aa) | This paper | N/A |
| pRK5-myc-Copine-6-Start-C2B (1-243aa) | This paper | N/A |
| pRK5-myc-Copine-6-LinkerB-End (306-557aa) | This paper | N/A |
| pX458-SpCas9(BB)-T2A-GFP | Addgene (F. Zhang) | Addgene plasmid<br>#48138 |
| pTrcHis-Copine-6<br>(wild-type, C2Amt and C2Bmt) | This paper | N/A |
| pX458-Copine-6-sgRNA2-Cas9-T2A-GFP | This paper | N/A |
| pX458-Copine-6-sgRNA3-Cas9-T2A-GFP | This paper | N/A |
| pU6-(BbsI)-CBh-Cas9-T2A-mCherry | Addgene (R. Kühn) | Addgene plasmid<br>#64324 |
| pU6-Copine-6-sgRNA3-Cas9-T2A-mCherry | This paper | N/A |
| **Software and Algorithms** |  |  |
| Fiji | http://fiji.sc | RRID:SCR_002285 |
| Image Studio Lite | https://www.licor.com/bio/image-studio-lite/ | RRID:SCR_015795 |
| GraphPad Prism 9 | https://www.graphpad.com/ | RRID:SCR_002798 |

### RESOURCE AVAILABILITY

#### Lead Contact

Further information and requests for resources and reagents should be directed to, and will be fulfilled by the Lead contact, Victor Anggono.

#### Materials Availability

All unique/stable reagents generated in this study are available from the Lead Contact upon request.

#### Data and Code Availability

This study did not generate datasets/code.

### EXPERIMENTAL MODEL AND SUBJECT DETAILS

#### Rats

Adult female Sprague-Dawley rats and their embryos (males and females, embryonic day 18) were used for the preparation of primary hippocampal neurons. All research procedures involving the use of animals were conducted in accordance with the Australian Code of Practice for the care and use of animals for scientific purposes and were approved by the University of Queensland Animal Ethics Committee (QBI/047/18 and 2021/AE511).

#### HEK293T cells

HEK293T cells were obtained from ATCC (CRL-3216). Cells were grown in DMEM with 4.5g/L glucose (GIBCO) supplemented with 10% FBS (Invitrogen) and 50 U/ml penicillin, 50 μg/ml streptomycin (GIBCO) in a humidified 5% CO_2_ incubator at 37°C.

### METHOD DETAILS

#### DNA constructs

Copine-6 cDNA was isolated from rat brain cDNA library by standard PCR with the following primers: 5’-GGGGATC<u>GTCGAC</u>CATGTCAGACCCAGAGATG-3’ (sense) and 5’-GGATT<u>GCGGCCGC</u>TCATGGGCTGGGGCTGGG-3’ (anti-sense), and cloned into the pRK5-myc vector using the *Sal*I and *Not*I restriction sites. Copine-6 Ca^2+^-binding mutants (C2Amt and C2Bmt) were generated by standard site-directed mutagenesis protocol. sgRNAs that target the rat copine-6 sequence 5’-CACCCGCAGCTCTACTCGAG-3’ (sgRNA2) and 5’-GAGCCTCTCGAGTAGAGCTG-3’ (sgRNA3) were cloned into pLentiCRISPR-GFP lentiviral vector (Incontro et al., 2018), pX458-Cas9-T2A-GFP and pU6-Cas9-T2A-mCherry. The previously described shRNAs targeting rat Syt1 5’-GAGCAAATCCAGAAAGTGCAA-3’ and rat Syt7 5’-GATCTACCTGTCCTGGAAGAG-3’ were cloned into the FG12 lentiviral vector (Wu et al., 2017).

#### Antibodies

The following antibodies were obtained commercially: rabbit anti-GluA1 C-terminal (ab1504, Millipore), rabbit anti-GluA1 N-terminal (abn241, Millipore), mouse anti-GluA1 C-terminal (182011, Synaptic Systems), mouse anti-β-actin sc-47778, Santa Cruz Biotechnology), rabbit anti-Copine-6 (13782-1-AP, ProteinTech), mouse anti-Copine-6 (sc-136357, Santa Cruz Biotechnology), chicken anti-MAP2 (ab92434, Abcam), mouse anti-myc (sc-40, Santa Cruz Biotechnology), mouse anti-GST (66001-2-Ig, ProteinTech), mouse anti-synaptotagmin-1 (105011, Synaptic Systems) and rabbit anti-synaptotagmin-7 (105173, Synaptic Systems).

Alexa-conjugated anti-rabbit secondary antibodies were purchased from Thermo Scientific, and Alexa-conjugated anti-chicken antibodies were bought from Abcam. HRP-conjugated secondary antibodies were obtained from GE Healthcare. Can Get Signal® Immunoreaction Enhancer Solution (Cat# NKB-601, Toyobo Life Science, Japan) was used to dilute rabbit anti-Copine-6 (ProteinTech) antibodies for immunofluorescence analysis.

#### Purification of recombinant Copine-6 protein

Plasmids encoding His-Copine-6 (WT, C2Amt or C2Bmt) were transformed into BL21(DE3) *E. coli* cells and cultured in LB broth at 37°C until the density reached an OD of 0.8. The expression of recombinant proteins was induced by adding 0.5 mM isopropyl 1-thio-β-D-galactopyranoside (IPTG) to the culture medium and the cultures were allowed to grow at 20°C overnight. Cells were harvested by centrifugation at 6,000 *g* for 10 min at 4°C. Cell pellets were resuspended in ice-cold buffer (50 mM Tris (pH 8.0), 300 mM NaCl, 50 μg/mL of benzamidine, 100 units DNaseI, 2 mM β-mercaptoethanol). The cells were lyzed by mechanical disruption at 30 kpsi using a Constant systems cell disrupter. The lysate was cleared by centrifugation at 50,000 *g* for 30 min at 4°C. Proteins were purified using affinity chromatography on a Ni-NTA (GE healthcare) gravity column. Proteins were eluted in 50 mM Tris (pH 8.0), 300 mM NaCl and 250 mM imidazole. The eluted affinity-purified proteins were then subjected to size exclusion chromatography using a Superdex-200 16/600 HiLoad column, pre-equilibrated with 50 mM Tris (pH 8.0), 100 mM NaCl and 2 mM DTT, and attached to an AKTA pure (GE Healthcare). The purified protein was concentrated to 10 mg/mL using a Centricon Ultra-10 kDa centrifugal filter (Millipore, USA) for isothermal titration calorimetry experiments. The concentration of protein was determined by the Bradford assay after each purification step.

#### Isothermal titration calorimetry

The affinities of His-Copine-6 WT, C2Amt or C2Bmt binding to Ca^2+^ were determined using a Microcal iTC200 instrument (Malvern, UK). Experiments were performed in 25 mM Tris (pH 8.0) and 100 mM NaCl. Two mM CaCl_2_ solution was titrated into 100 µM of purified His-Copine-6 in 13 × 3.22 µL aliquots at 25°C. The dissociation constants (*K*_d_), enthalpy of binding (ι1*H*) and stoichiometries (N) were obtained after fitting the integrated and normalized data to a single-site binding model. The stoichiometry was refined initially, and the value obtained was close to 1; then, N was set to 1.0 for calculation. The apparent binding free energy (ι1*G*) and entropy (ι1*S*) were calculated from the relationships ι1*G* = RTln(*K*_d_) and ι1*G* = ι1*H* - Tι1*S*. All experiments were performed at least in duplicate to check for the reproducibility of the data.

#### Subcellular fractionation and GST pull-down assays in HEK293T cells

HEK293T cells were grown in DMEM with 4.5 g/L glucose supplemented with 10% FBS, 50 U/mL penicillin and 50 mg/mL streptomycin in a humidified 5% CO_2_ incubator at 37°C. A subcellular fractionation assay was performed to analyze the association of Copine-6 with cell membranes. Briefly, HEK293T cells were transfected with either pHA-N1-Copine-6 WT, C2Amt or C2Bmt for 48 h. Cells were then treated with either 2 mM EGTA or 1 mM CaCl_2_ + 2 μM ionomycin for 5 min. Cells were lyzed in ice-cold lysis buffer (5 mM NaHCO_3_, 1 mM KH_2_PO_4_, 0.5 mM MgCl_2_, 10 mM NaCl, 10 mM HEPES pH 7.5) containing EDTA-free protease inhibitor cocktail (Roche). Homogenization was then performed, and isotonic conditions were restored by adding sucrose to a final concentration of 227 mM. Nuclei and cell debris were removed by centrifugation at 6,300 *g*, 4°C for 5 min. Cytoplasmic and membrane fractions were obtained by ultracentrifugation at 100,000 *g*, at 4°C for 30 min.

GST pull-down assays were performed to analyze the interaction between Copine-6 and C-terminal tails of GluA1 or GluA2. HEK293T cells were co-transfected with either pCIS-empty (GST), pCIS-GluA1 C-tails (either full-length or truncated) or pCIS-GluA2 C-tail, together with pRK5-myc-Copine-6 (either full-length or truncated) for 24 h. Cells were lyzed in ice-cold cell lysis buffer (1% Triton X-100, 1 mM EDTA, 1 mM EGTA, 50 mM NaF, 5 mM Na-pyrophosphate in PBS) supplemented with EDTA-free protease inhibitor cocktail (Roche). Lysates were cleared by centrifugation at 20,627 *g*, 4°C for 20 min. A small amount of cleared lysate was collected for input and the remaining lysate was incubated with glutathione agarose beads (Thermo Scientific) overnight at 4°C. Beads were washed three times with ice-cold cell lysis buffer and bound proteins were eluted with 2X SDS sample buffer at 100°C for 10 min. Samples from GST pull-down and subcellular fractionation assays were analyzed by Western blotting.

#### Rat primary neuronal culture and transient transfection

Primary rat hippocampal and cortical neurons were prepared from Sprague-Dawley rat embryos at embryonic day 18 as previously described (Widagdo et al., 2015). Briefly, isolated cortical and hippocampal tissues were dissociated using 30U of papain solution (Worthington) for 20 min in a 37°C water bath. Tissues were then triturated using fire-polished glass Pasteur pipettes to generate a single-cell suspension. Neurons were plated on a poly-L-lysine-coated 12-well plate or 6-cm dish at a density of 8 x 10^4^ cells (hippocampal neurons, 12-well plate), 2 x 10^5^ cells (cortical neurons, 12-well plate) or 1 x 10^6^ cells (cortical neurons, 6-cm dish). Neurons were cultured in Neurobasal growth medium supplemented with 2% B-27, 2 mM GlutaMAX and 1% penicillin/streptomycin and incubated in a 37°C humidified tissue culture incubator with 5% CO_2_. Neurons were fed twice a week with Neurobasal medium containing 0% and 1% FBS for hippocampal and cortical neurons, respectively. To stop glial proliferation in cortical cultures, 5 μM uridine (Sigma) and 5 μM 5’-fluoro-2’-deoxyuridine (Sigma) were added to the culture medium at days *in vitro* (DIV) 5. Hippocampal neurons were transfected at DIV 12-13 using Lipofectamine 2000 (Invitrogen) according to the manufacturer’s protocol and processed at DIV 15-16.

#### Lentivirus packaging and transduction

Lentiviral particles were generated in HEK293T cells that had been transfected with 7 μg of the plasmid of interest, 3 μg each of pMD2.G envelope plasmid, pRSV-Rev encoding plasmid and pMDLg/pRRE packaging constructs via the calcium-phosphate precipitation method. Lentivirus-containing supernatant was collected 48 h after transfection and filtered through a 0.45 μm low protein-binding cellulose acetate membrane. Lentiviral particles were pelleted either by ultracentrifugation at 106,559 *g* on a Beckman SW 32 Ti rotor for 2 h at 4°C or using the PEG-it Virus Precipitation Solution (System Biosciences) following the manufacturer’s protocol. The viral concentrate was resuspended in a Neurobasal medium, snap-frozen in liquid nitrogen and stored at −80°C. Cortical neurons were transduced with lentiviral particles at DIV 8-9 (overnight) and DIV 12 (6 h) for Copine-6 sgRNA-knockout and overexpression, respectively. Transduced neuronal cultures were incubated for a further 3-6 days prior to processing.

#### Glycine-induced chemical LTP (chem-LTP) and tetrodotoxin (TTX) treatment

Chem-LTP was induced on mature primary neurons at DIV 15-16. Briefly, neurons were washed and incubated in pre-warmed artificial cerebrospinal fluid with low Mg^2+^ (ACSF; containing 0.4 mM MgCl_2_, 2 mM CaCl_2_, 120 mM NaCl, 5 mM KCl, 30 mM glucose, 25mM HEPES, pH 7.4) supplemented with 20 µM bicuculline (Abcam), 5 µM strychnine (Sigma) and 0.5 µM TTX (Abcam) for 1 h at 37°C. Induction of chem-LTP was carried out by incubating neurons in pre-warmed low Mg^2+^ ACSF containing 200 µM glycine, 20 µM bicuculline and 5 µM strychnine for 5 min at room temperature. To induce homeostatic synaptic up-scaling of AMPARs, primary cortical neurons were treated with 2 µM TTX (Abcam) for 48 h at DIV10-12.

#### Co-immunoprecipitation assay

Primary rat cortical neurons (DIV 22) were lyzed in ice-cold cell lysis buffer containing EDTA-free protease and phosphatase inhibitor cocktails for 30 min at 4°C and cleared by centrifugation at 20,627 *g* for 20 min. Lysates were pre-cleared with 25 μL of Protein G agarose beads (GE Healthcare). At the same time, 25 μL of Protein G agarose beads were pre-coupled with either 4 μg of mouse anti-Copine-6 antibodies, 5 μg of anti-GluA1 antibodies or 5 μg of normal mouse IgG for 1h at 4°C. Pre-cleared lysates were subjected to immunoprecipitation at 4°C overnight. Beads were washed three times, and eluted with 2X SDS sample buffer, followed by Western blotting analysis.

#### Copine-6 translocation assay

To analyze the activity-dependent translocation of Copine-6 in dendrites, primary hippocampal neurons were co-transfected with pEGFP-N1-Copine-6 (WT, C2Amt or C2Bmt) and a cytosolic marker td-Tomato at DIV 13. At DIV 15, the translocation of Copine-6-GFP in live neurons was visualized before and after 3 min treatment with 10 μM NMDA + 100 μM glycine. Images were taken every 5 s over 400 s using the Yokogawa spinning disk confocal microscope. The translocation index of Copine-6-GFP was calculated by dividing the membrane to cytosol fluorescence intensity ratio in the NMDA-treated vs non-stimulated conditions, normalized by changes in the td-Tomato fluorescence intensities.

#### Immunolabelling of surface GluA1

An antibody-feeding assay was performed to determine the level of AMPARs on the plasma membrane. Primary hippocampal neurons were washed twice in ice-cold ACSF, followed by incubation with rabbit anti-GluA1 N-terminal antibodies (1:100) for 30 min at 4°C. Neurons were subsequently fixed with Parafix solution (4% paraformaldehyde, 4% sucrose in PBS) for 10 min, blocked (10% normal goat serum in PBS) for 1 h, and incubated with Alexa-568-conjugated anti-rabbit IgG secondary antibodies (1:100) for 1 h. After washing, neurons were again fixed for 2 min, permeabilized (0.125% Triton X-100 in PBS) for 10 min and blocked for 1 h. Total GluA1 was then labeled with rabbit anti-GluA1 C-terminal antibodies (1:500) overnight at 4°C and visualized by staining with Alexa-647-conjugated anti-rabbit IgG antibodies for 1 h. Images were collected with a 63X oil-immersion objective on a Zeiss LSM510 confocal microscope. Fluorescence intensities of surface and total GluA1 were quantified using ImageJ software (National Institutes of Health).

#### SEP-GluA1 insertion assay

Primary hippocampal neurons were transfected with super-ecliptic pH-sensitive GFP-tagged GluA1 (SEP-GluA1) between DIV 11 and 12, and imaged 3 days later in ACSF (pH 7.4) with or without glycine stimulation. Images were collected with a 100X oil-immersion objective on a Zeiss ELYRA microscope using the TIRF mode. To visualize newly inserted AMPA receptors on the dendritic membranes, pre-existing surface SEP-GluA1 was photobleached with 100% laser power for 10 sec before data acquisition. Live cell images were captured at 0.5 Hz for 5 min. Signals that last for at least four frames (8 sec) were manually scored as insertion events from the *y*-t rendered images. The *y*-t rendering was performed on Image J software as described previously (Lin et al., 2009). Data were expressed as total number of GluA1 insertions per μm of dendrite over 5 min. The typical length of secondary dendritic segments analyzed was 37-45 μm.

#### Surface biotinylation assay

The surface biotinylation assay was carried out to measure the levels of endogenous GluA1 on the plasma membrane. Live neurons were rinsed with cold ACSF (pH 8.2) and incubated with ACSF (pH 8.2) containing 0.5 mg/ml sulfo-NHS-SS-Biotin (CovaChem) for 30 min at 4°C. Free biotin was quenched by washing cells three times with ice-cold Tris-buffered saline (TBS). Neurons were then lyzed with RIPA buffer (1% Triton X-100, 0.5% Na-deoxycholate, 0.1% SDS, 2 mM EDTA, 2 mM EGTA, 50 mM NaF, 10 mM Na-pyrophosphate, 150 mM NaCl, 50 mM Tris pH 7.4) supplemented with EDTA-free protease inhibitor cocktail for 30 min at 4°C. Lysates were cleared by centrifugation at 20,627 *g* at 4°C for 20 min. A small fraction of cleared lysates were collected as input to determine the total GluA1 level, and the remaining cleared lysates were incubated with Neutravidin beads (Thermo Scientific) overnight at 4°C to isolate surface biotinylated proteins. The beads were washed three times with ice-cold RIPA buffer and bound proteins were eluted with 2X SDS sample buffer and heated at 50°C for 30 min. Samples were analyzed by Western blotting. The ratio of surface/total GluA1 of each sample was normalized to control.

#### Western blotting

Samples were loaded in 7.5 or 12% SDS-PAGE gels and separated at 110 V for 1-2 h. Proteins were then transferred to a PVDF membrane at 100 V for 2 h. Membranes were blocked in 5% skim milk (in TBS containing 0.1% Tween-20, TBS-T) for 1 h and washed in TBS-T three times at 5 min intervals prior to overnight incubation with primary antibodies at 4°C. Membranes were washed in 1% milk/TBS-T five times and incubated with HRP-conjugated secondary antibodies (GE Healthcare, 1:10,000) for 1 h at room temperature. They were washed extensively and developed using the enhanced chemiluminescent method (PerkinElmer). Images were acquired with a LiCOR imaging system and quantified with the ImageStudio software.

### QUANTIFICATION AND STATISTICAL ANALYSIS

The sample size (*n*) reported in figure legends represents individual neurons or wells generated from at least three independent experiments, unless otherwise stated. Statistical analysis was performed in Graph Pad Prism 9.0 using one-way analysis of variance (ANOVA) with Tukey’s or Sidak’s post-hoc multiple comparisons tests. For comparison between two groups, a two-tailed Mann-Whitney’s test was employed. All data are reported as mean ± standard error of the mean (SEM).

---

**Figure S1 (related to Figure 3).**
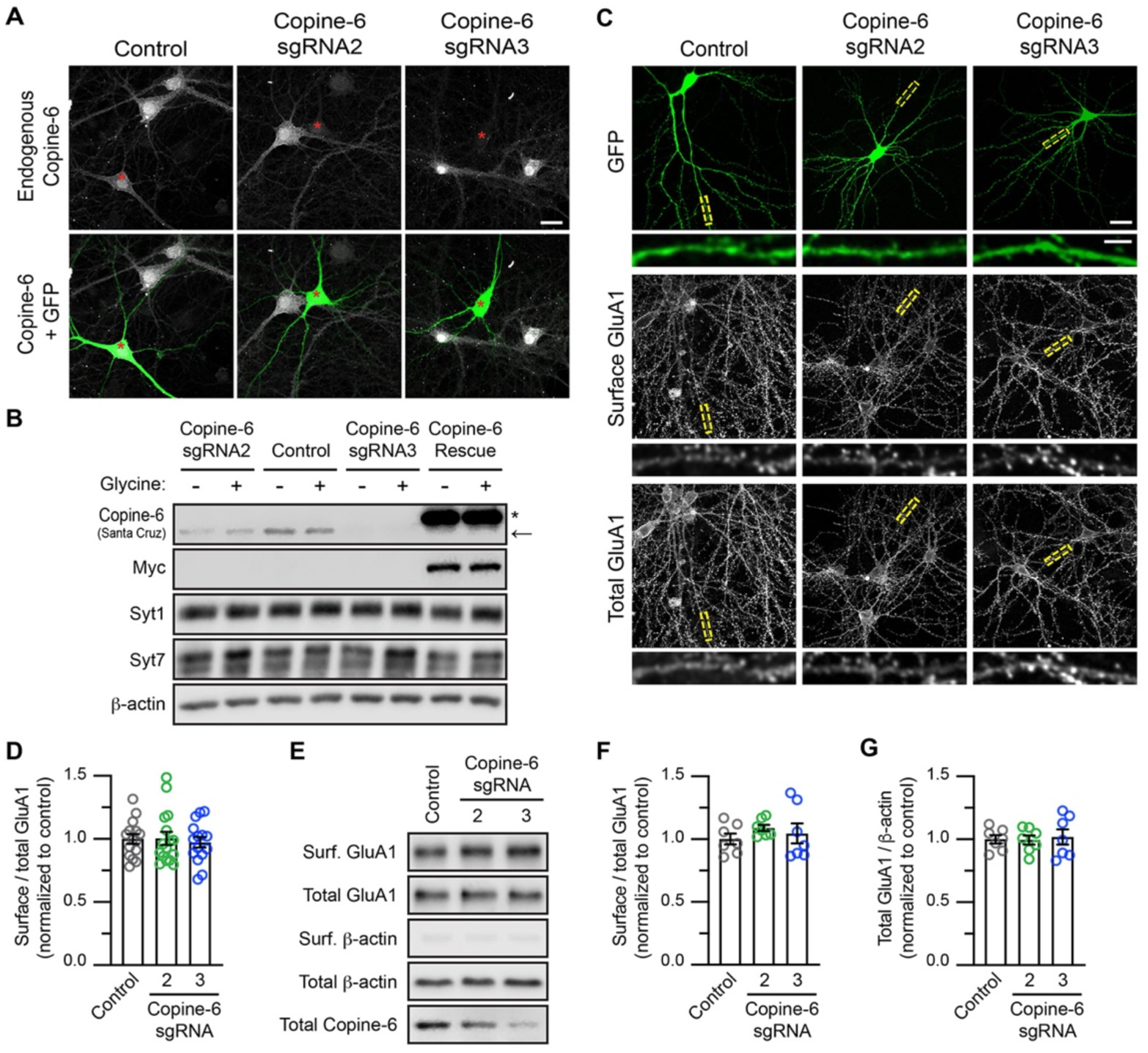
Loss of Copine-6 expression does not affect the steady-state expression of AMPARs under basal conditions. (A) Knockdown efficiency of endogenous Copine-6 in primary neurons using the CRISPR-Cas9 sgRNA system. Primary hippocampal neurons were transfected with plasmids encoding Cas9-GFP, either alone or with Copine-6-sgRNA2 or sgRNA3 at DIV12. Neurons were fixed, permeabilized and stained with specific antibodies against Copine-6 (gray). Transfected neurons (green) are marked with red asterisks. Scale bar, 20 μm. (B) Copine-6 depletion did not alter the levels of Syt-1 or Syt-7 proteins. Primary cortical neurons were transduced with lentiviral particles expressing Cas9-GFP with or without Copine-6 sgRNA2 or sgRNA3 at DIV 9. For the Copine-6 rescue group, neurons were further transduced with lentiviral particles expressing sgRNA3-resistant Myc-Copine-6 WT at DIV 12. Lysates from DIV 15 neurons were resolved by SDS-PAGE and analyzed by Western blotting with specific antibodies against Copine-6, Syt-1, Syt-7, β-actin and Myc. Endogenous and exogenous Copine-6 protein bands are indicated by an arrow and asterisk, respectively. (C) Hippocampal neurons expressing Cas9-GFP, either alone or with Copine-6 sgRNA2 or sgRNA3 were incubated with a specific antibody against the N-terminus of GluA1 at 4°C (surface GluA1), followed by fixation, permeabilization and staining with a specific antibody against the C-terminus of GluA1 (total GluA1). Representative images of the surface and total GluA1 staining in neurons from each group are shown (top panels). Scale bar, 20 μm. The bottom panels show enlarged images of dendritic segments within the boxed regions. Scale bar, 5 μm. (D) Quantification of the surface/total GluA1 ratio normalized to control neurons (*n* = 15-16 neurons per group from 3 independent experiments). (E) Primary cortical neurons were transduced with lentiviral particles expressing Cas9-GFP, either alone or with Copine-6 sgRNA2 or sgRNA3 at DIV 9. Transduced neurons were subjected to a surface biotinylation assay at DIV 15. The relative amounts of surface and total proteins were assessed by Western blotting using specific antibodies against GluA1, Copine-6 and β-actin. (F and G) Quantification of the surface/total GluA1 (F) and the total GluA1/ β-actin (G) ratios in Copine-6 depleted neurons, normalized to control neurons (*n* = 7 cultures per group from 3 independent experiments). All data represent mean ± SEM.

**Figure S2 (related to Figure 3).**
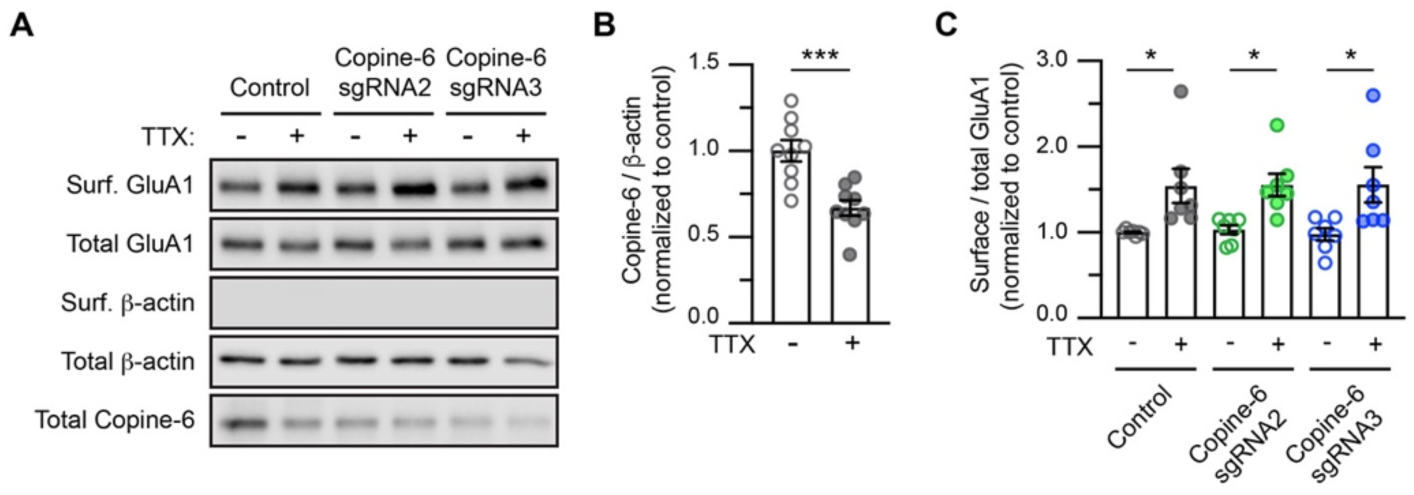
Copine-6 is not required for TTX-induced homeostatic synaptic up-scaling of AMPARs. (A) Primary cortical neurons were transduced with lentiviral particles expressing Cas9-GFP, either alone or with Copine-6 sgRNA2 or sgRNA3 at DIV 7. At DIV 10, transduced neurons were treated with 2 μM TTX for 48 h and subjected to a surface biotinylation assay at DIV 12. The relative amounts of surface and total proteins were assessed by Western blotting using specific antibodies against GluA1, Copine-6 and β-actin. (B) Chronic neuronal inactivity significantly reduces Copine-6 protein expression (*n* = 9 cultures from 5 independent experiments). \*\*\**P* < 0.001 using a Mann-Whitney t-test. (C) Quantification of the surface/total GluA1 ratio (*n* = 7-9 cultures from 5 independent experiments). \**P* < 0.05 using one-way ANOVA with Sidak’s multiple comparison test.

**Figure S3 (related to Figure 4).**
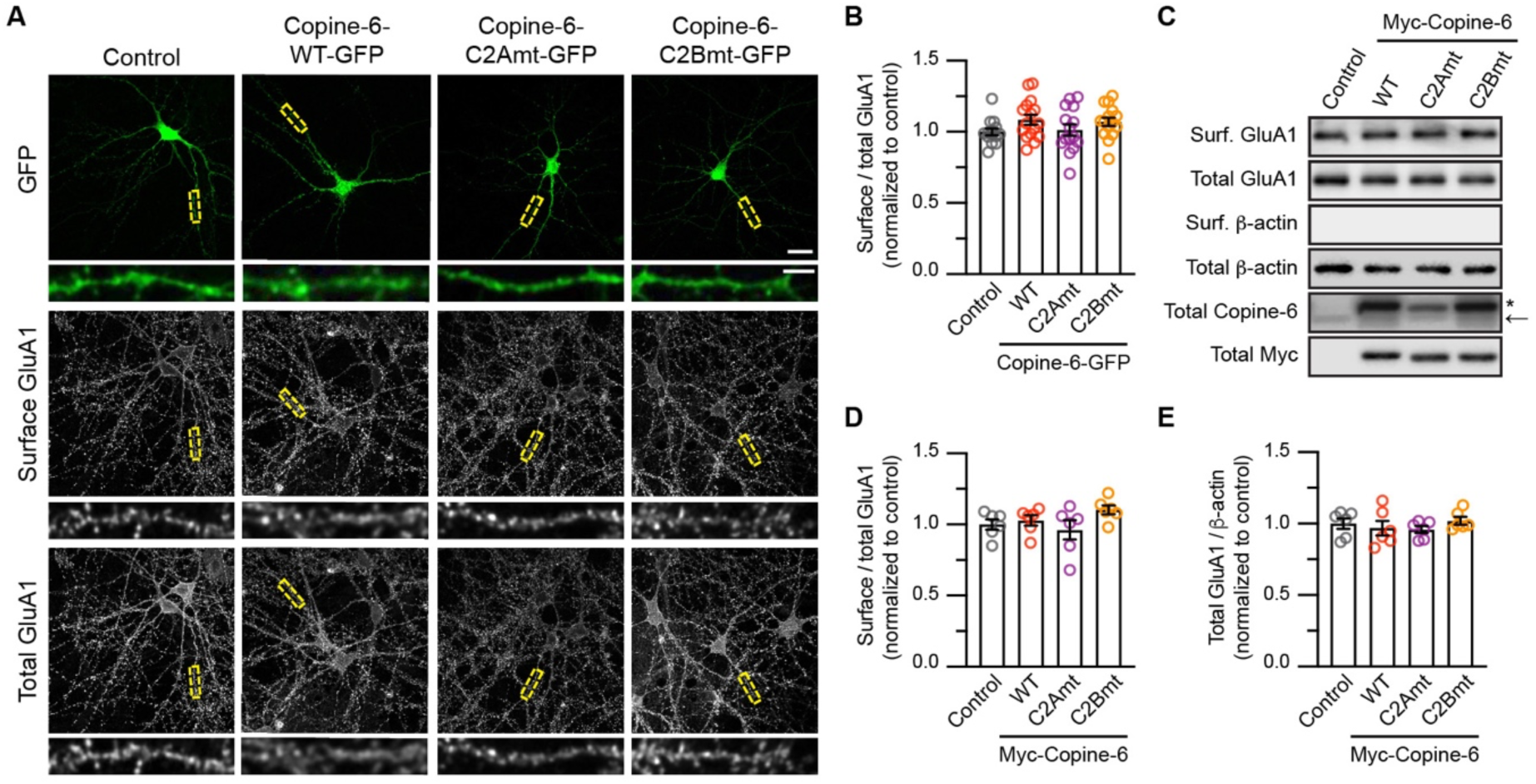
Overexpression of Copine-6 Ca2+-binding mutants does not affect the steady-state levels of surface AMPARs under basal conditions. (A) Hippocampal neurons expressing GFP alone (control) or Copine-6-GFP (WT, C2Amt or C2Bmt) were incubated with a specific antibody against the N-terminus of GluA1 at 4°C (surface GluA1), followed by fixation, permeabilization and staining with a specific antibody against the C-terminus of GluA1 (total GluA1). Representative images of the surface and total GluA1 staining in neurons from each group are shown (top panels). Scale bar, 20 μm. The bottom panels show enlarged images of dendritic segments within the boxed regions. Scale bar, 5 μm. (B) Quantification of the surface/total GluA1 ratio normalized to control neurons (*n* = 14-16 neurons per group from 3 independent experiments). (C) Primary cortical neurons were transduced with lentiviral particles expressing GFP alone (control) or Myc-Copine-6 (WT, C2Amt or C2Bmt) at DIV 12. Transduced neurons were subjected to a surface biotinylation assay at DIV 15. The relative amounts of surface and total proteins were assessed by Western blotting using specific antibodies against GluA1, Copine-6, Myc and β-actin. (D and E) Quantification of the surface/total GluA1 (D) and the total GluA1/ β-actin (E) ratios, normalized to control neurons (*n* = 6 cultures per group from 3 independent experiments). All data represent mean ± SEM.

**Figure S4 (related to Figure 4).**
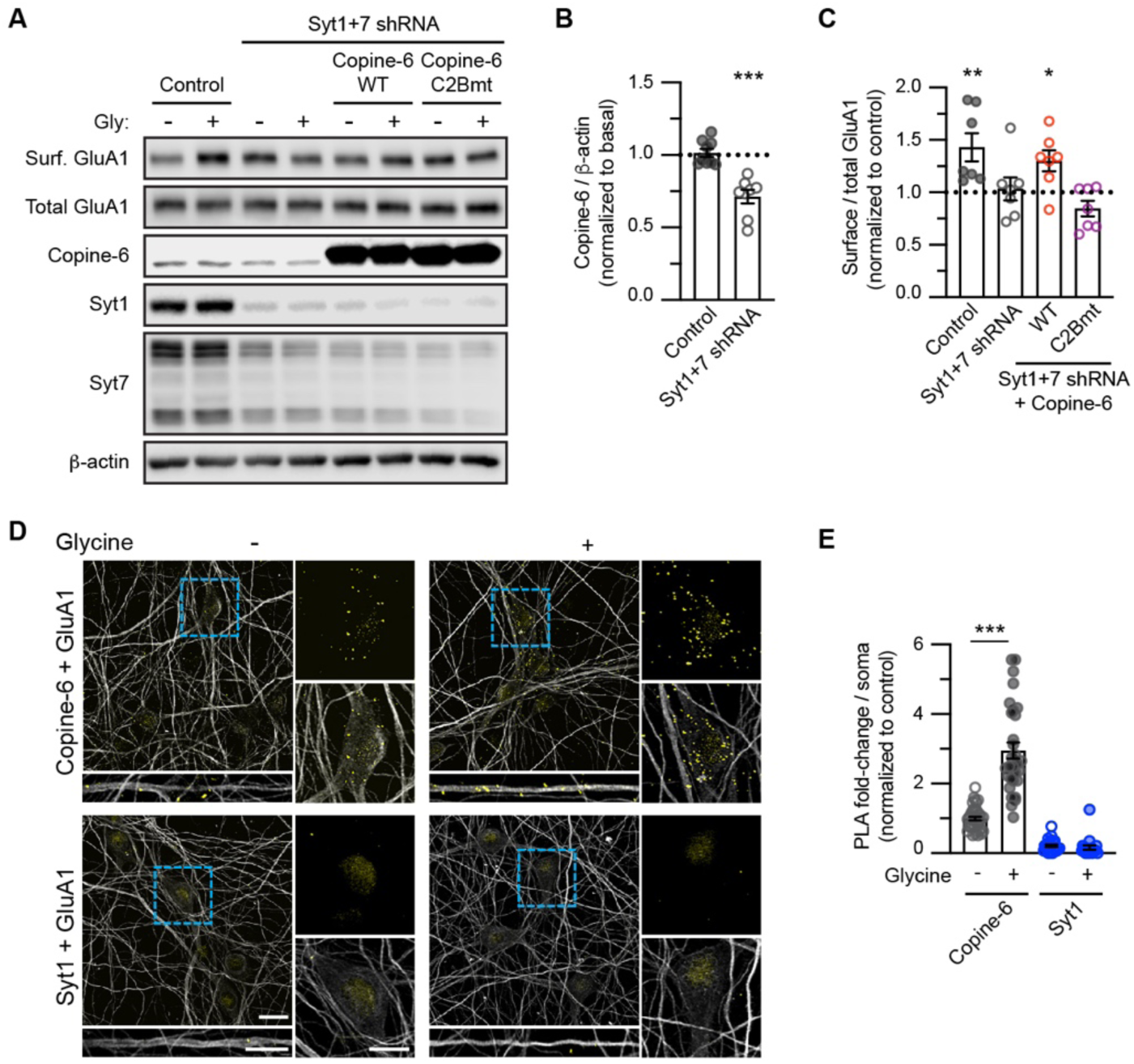
Overexpressing Copine-6 WT restores glycine-induced insertion of AMPARs to the plasma membrane in Syt-1/Syt-7 depleted neurons. (A) Primary cortical neurons were transduced with lentiviral particles expressing GFP alone (control) or Syt1+7 shRNAs plus GFP at DIV9. At DIV 12, some of the Syt1+7 depleted neurons were further transduced with lentiviral particles expressing Myc-Copine-6, either WT or the C2Bmt. Neurons were stimulated with glycine for 5 min and subjected to a surface biotinylation assay at DIV15. The relative amounts of surface and total proteins were assessed by immunoblotting with specific antibodies against GluA1, copine-6, Syt1, Syt7 and β-actin. (B) Loss of Syt-1 and Syt-7 significantly reduces Copine-6 protein expression (*n* = 7 cultures from 7 independent experiments). \*\*\**P* < 0.001 using a Mann-Whitney t-test. (C) Quantification of the surface/total GluA1 ratios, normalized to unstimulated groups (*n* = 7 cultures per group from 7 independent experiments). \**P* < 0.05; \*\**P* < 0.01 using one-way ANOVA with Sidak’s multiple comparison test. All data represent mean ± SEM. (D) Primary hippocampal neurons (DIV 15) were stimulated with glycine for 5 min and subjected to a proximity ligation assay (PLA) with GluA1 and Copine-6 (upper panels) or GluA1 and Syt-1 (bottom panels) antibodies. Representative images showing the PLA signals (yellow) and MAP2 staining (gray) in the soma or dendrites (insets). Scale bars, 10 μm. (E) Quantification of the number of PLA puncta in the soma. Data are normalized to the unstimulated GluA1-Copine-6 group (*n* = 23-40 neurons per group from 2 independent experiments). All data are represented as mean ± SEM. \*\*\**P* < 0.001 using one-way ANOVA with a Tukey’s multiple comparison test.

**Table S1 (related to Figure 1).** Thermodynamic parameters of Copine-6 binding to Ca^2+^. 2 mM CaCl_2_ was titrated into 100 μM of recombinant His-Copine-6 full-length proteins, either wild-type (WT), C2Amt (D93N/E95Q) or C2Bmt (D229N/D231N/D237N) Ca^2+^-binding mutants at 25°C. Data represent mean ± SEM of 3 independent experiments. NBD: no binding detectable.

| Protein + $\text{Ca}^{2+}$ | n<br>(stoichiometry) | Kd ($\mu\text{M}$) | $\Delta\text{H}$ (kcal/mol) | T$\Delta\text{S}$ (kcal/mol) | $\Delta\text{G}$ (kcal/mol) |
| --- | --- | --- | --- | --- | --- |
| Copine-6 WT | 1.23 ± 0.05 | 9.3 ± 1.8 | -1.17 ± 0.42 | 5.69 ± 0.23 | -6.86 ± 0.73 |
| Copine-6<br>C2Amt | 0.97 ± 0.02 | 30.2 ± 3.06 | -2.12 ± 0.28 | 3.96 ± 0.57 | -6.08 ± 0.89 |
| Copine-6<br>C2Bmt | NBD | NBD | NBD | NBD | NBD |

